# Are island plants really poorly defended? No support for the loss of defense hypothesis in domatia-bearing plants

**DOI:** 10.1101/2021.11.11.468197

**Authors:** M. Biddick

## Abstract

The loss of defense hypothesis posits that island colonizers experience a release from predation on the mainland and subsequently lose their defensive adaptations. While support for the hypothesis is abundant, it has never been tested in domatia-bearing plants. Leaf domatia are cave-like structures produced on the underside of leaves that facilitate a defensive mutualism with predatory and fungivorous mites. I tested the loss of defense hypothesis in six domatia-bearing taxa inhabiting New Zealand and its offshore islands. No support for the loss of defense hypothesis was found. Changes in domatia investment were instead associated with changes in leaf size – a trait that has been repeatedly observed to undergo rapid evolution on islands. Overall results demonstrate that not all types of defense are lost on islands, suggesting a higher-resolution approach is needed when studying the evolution of defense on islands.

## 2. Introduction

Island organisms can often differ from their mainland counterparts in remarkable yet predictable ways. Collectively, these ecological, morphological, and behavioural differences comprise what is known as the *‘island syndrome’* [1–3]. One of the most remarkable trends comprising the island syndrome is the loss of defensive adaptations. The loss of defense hypothesis posits that island organisms lose defensive adaptations after colonizing isolated islands because they no longer provide a fitness benefit in the absence of mainland predators [4]. For instance, many endemic island birds like the Dodo of Mauritius exhibit no fear of mammalian predators – including humans (unfortunately to their detriment). In fact, almost half of the endemic avifauna of the New Zealand archipelago went extinct following the introduction of mammalian predators [5, 6]. Similar behavioural shifts are seen in other prey taxa like insular foxes, wallabies, and lizards [7, 8]. However, defensive adaptations are not only lost in island animals.

Many island plants have also lost defensive adaptations [2, 9, 10]. While prey animals change in response to the absence of mainland predators in their new environment, plants change in response to absent mainland herbivores [11]. Leaf spines, for example, are effective physical deterrents of grazing ungulates like deer and cattle [12, 13]. However, on the California Channel Islands where ungulates have historically been absent (until recent introductions), many endemic plants produce larger leaves, fewer spines, and less phenol compounds than their most closely related mainland sister taxa [14]. Another remarkable loss of defense is seen in *Pseudopanax crassifolius* (lancewood) from New Zealand. Lancewood saplings produce long, hardened leaves with lateral leaf spines that are advertised with aposematic colouration [15]. Thought to deter large (now extinct) browsing birds called *Moa* [16], these morphological adaptations have been lost in populations of *P. crassifolius* inhabiting the Chatham Islands some 800km away (where *Moa* were absent) [17]. However, not all forms of plant defense are physical.

Some plants produce cave-like structures on the underside of their leaves known as domatia that facilitate a defensive mutualism with predatory and fungivorous mites [18, 19]. Leaf domatia provide a refuge for oviposition and shelter from the environmental extremes of leaf surfaces, which mites in turn protect via the consumption of herbivorous arthropods and fungal pathogens [20]. Some 2000 taxa of domatia-bearing plants belonging to 227 plant families have been described [21]. Intriguingly, no study has documented whether this defensive mutualism is lost on islands. To test this formally, I quantified domatia production in six cosmopolitan taxa inhabiting the New Zealand mainland and its surrounding offshore islands (figure 1).

**Figure 1.**
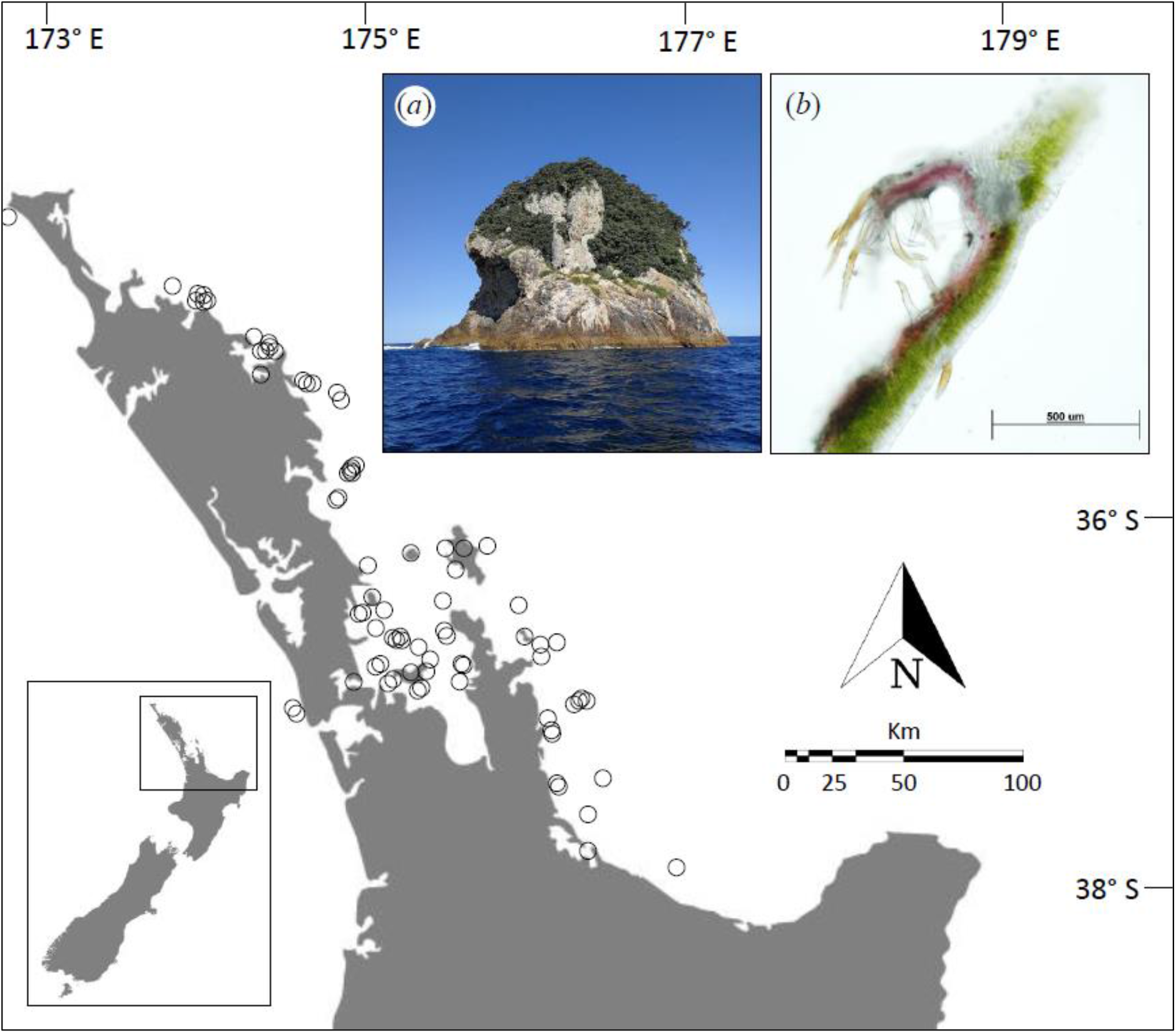
Study area. (*a*) Remote islands off the east coast of the New Zealand ‘mainland’ were used to test the loss of defense hypothesis. (*b*) Leaf domatia are cave-like structures produced on the underside of leaves that facilitate a defensive mutualism with predacious and fungivorous mites (example of tuft domatia in *Carpodetus serratus* shown, photo credit: Morgan Ngata). Open circles denote individual offshore islands.

## 3. Methods

### (a) Data collection

Data collection was conducted between June 2018 and April 2019. Individuals were chosen haphazardly while walking through easily accessible forest sections. Only fully expanded, mature leaves were measured. Leaf length was measured as the longest linear distance from the most proximal to the most distal point of the leaf lamina using a digital calliper. Leaf width was measured as the widest distance across the leaf lamina perpendicular to the leaf length measurement. Leaf area was calculated as the product of leaf length and leaf width. Although more accurate methods of estimating leaf area exist [e.g., using leaf shape correction factors or leaf scanners, 22]), leaf *x* width calculations sufficed for the purpose of this analysis as it is not concerned with among species differences in leaf size. Couched in other terms, any errors associated with leaf area estimates are consistent between island-mainland comparisons. Further, many of the islands included in this study are protected by the Department of Conservation, who maintain strict standards about the type of sampling that can be conducted on them. Domatia were then counted either by eye or USB microscope (for smaller domatia).

To expand upon data gathered in the field, pressed herbarium specimens from the Auckland War Memorial Museum Herbarium were measured. The Herbarium houses an extensive collection of high quality, preserved specimens of both indigenous and exotic plants spanning the full geographic extent of the New Zealand landmass – including its offshore islands. To standardize our sampling across specimens, 3 leaves per specimen were measured using the same methodology outlined above. Leaves were chosen haphazardly, and care was taken to not damage specimens (i.e., gloves and minimal calliper–specimen contact). Mainland sampling was conducted in the Kaimai-Mamaku Forest Park in Tauranga (37°41’S, 175°45’E). This site was chosen because the Kaimai Ranges span a large latitudinal extent of the north-eastern corner of New Zealand, and therefore represent the probable source pool for many island populations.

### (b) Data analysis

To test for mainland-island differences in domatia production, I conducted species-wise Welch *t*-tests. Data were left untransformed as the Welch *t*-test is robust to violations of unequal variance. The same approach was used to test for mainland-island differences in leaf size. All analyses and data visualization were conducted in the R environment (v. 3.0.1) using the ‘tidyverse’ collection of packages [23].

## 4. Results

No evidence for the loss of defense hypothesis was observed, with 2 taxa exhibiting marginally reduced domatia production under insular conditions, 2 taxa exhibiting no change, and 2 taxa showing increased domatia production (table 1, figure 2). Island populations of *C. rhamnoides* exhibited marginally higher rates of domatia production relative to mainland populations. Changes in domatia production were instead associated with changes in leaf size on islands (table 1).

**Table 1.** Results of species-wise Welch t-tests analysing mainland-island population differences in leaf size and domatia production. Samples sizes are denoted in parentheses for mainland and island populations, respectively. Significant *p*-values ≤ 0.05 are denoted in italics.

| species | leaf size |  |  | leaf domatia |  |  |
| --- | --- | --- | --- | --- | --- | --- |
| | $T$ | $p$ | insular change | $T$ | $p$ | insular change |
| *Coprosma repens* (30, 332) | -6.81 | $< 0.01$ | decrease | -2.44 | $0.02$ | decrease |
| *Coprosma rhamnoides* (64, 204) | 9.77 | $< 0.01$ | increase | 1.93 | $0.05$ | increase |
| *Coprosma robusta* (52, 60) | 2.34 | $0.02$ | increase | -1.49 | 0.14 | no change |
| *Coprosma lucida* (62, 245) | 4.42 | $< 0.01$ | increase | 2.56 | $0.01$ | increase |
| *Elaeocarpus dentatus* (80, 89) | 0.88 | 0.38 | no change | -2.49 | $0.02$ | reduce |
| *Vitex lucens* (30, 36) | 4.43 | $< 0.01$ | increase | 0.82 | 0.42 | no change |

**Figure 2.**
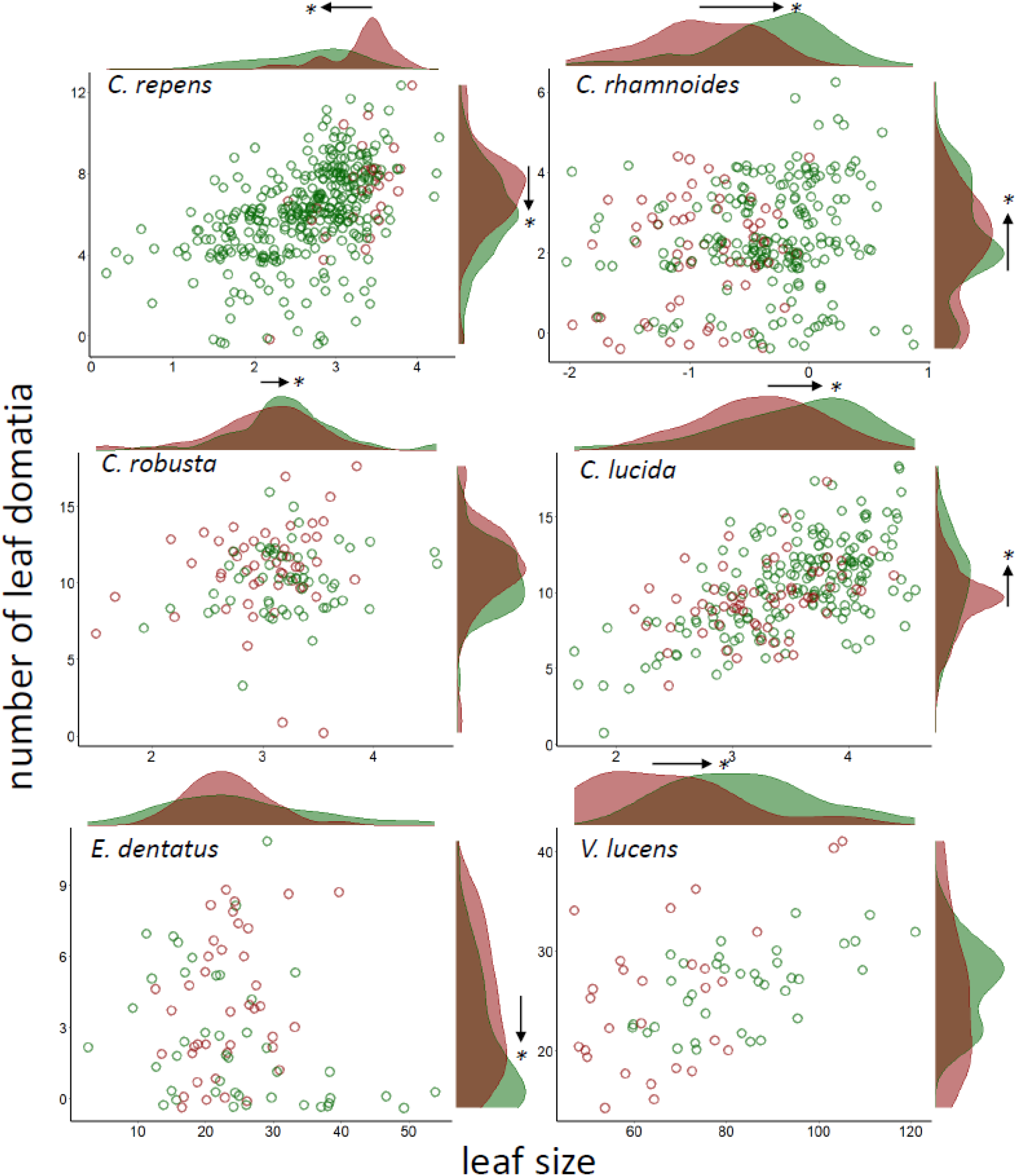
Leaf domatia production (*y*–axis) and leaf size (*x*–axis, logarithm-transformed) in six cosmopolitan plant taxa inhabiting the New Zealand mainland (red) and its offshore islands (green). Black arrows and asterisks denote significance and direction of change in each trait on islands.

## 5. Discussion

Many previous studies have documented the loss of defense adaptations in island organisms, and loss of defense is considered more generally to constitute part of the ‘island syndrome’ [1–3]. However, findings from this study demonstrate that not all types of defense are lost on islands. Only a marginal reduction in leaf domatia production was observed in two of the six taxa considered. The remaining taxa either showed no change or, contrastingly, increased domatia production on islands.

Why aren’t leaf domatia lost on islands? The putative explanation for defense loss in island organisms is the absence of mainland predators. For instance, mammals are typically poor dispersers and thereby lack the ability to colonize remote islands. Plants that have historically been defended against such mammalian predators, and subsequently colonize isolated islands, are freed from this predation pressure, and no longer benefit from the production of physical deterrents like thorns, spines, and prickles. The mites inhabiting leaf domatia, on the other hand, protect plants against insect and fungal attack – both of which disperse readily to islands [24].

Another potential explanation is that the islands considered in this study, and the flora inhabiting them, are too young to exhibit the loss of defense phenomenon. Indeed, previous work has demonstrated the effect of island age and time since divergence on phylogeographic patterns [25, 26]. Most of the islands considered here are continental break offs from the larger New Zealand landmass. As such, they are relatively young in comparison to true oceanic islands like Hawaii or Mauritius. This could explain why a marginal decrease in domatia production was observed in some taxa. Further, many islands are only weakly isolated from the New Zealand mainland, such that gene flow between island and mainland populations may exist, dampening the signal of an otherwise real effect [see 27 for discusssion].

Contrary to the loss of defense hypothesis, changes in domatia production were instead associated with changes in leaf size on islands. Prior work has demonstrated that leaf size in woody plants obeys the island rule [28] – a ubiquitous pattern in island evolution whereby species converge on intermediate sizes on islands. Changes in domatia production in island populations may therefore be explained by changes in leaf size if both traits are allometrically related (though allometrically-linked traits have been shown to evolve independently [29]).

In conclusion, island plants are not ‘poorly defended’ *per se*. Rather, they tend to lose adaptations to predators and competitors that are absent on the isolated islands they colonize. Because insect and fungal attack are still very much a threat on islands, the mutualistic relationship of plants with predacious and fungivorous mites facilitated by leaf domatia remains advantageous. Future work should test whether other types of defensive mutualisms are lost or reduced on islands, such as those with ants. Finally, the loss of defense hypothesis should be amended to reflect *changes* in defense rather than its loss. While island endemics are often poorly defended against introduced mammalian predators, many are exceptionally well protected against (now extinct) avian predators [30].

## Data accessibility

Data related to this article can be found at doi:10.5061/dryad.rv15dv48q.

## Author contributions

M.B collected the data, conducted analyses, and wrote the manuscript.

## Acknowledgements

I am indebted to KC Burns for his mentorship early in my academic career. I thank Ewen Cameron and all staff at the Auckland War Memorial Museum for providing access to the remarkable Herbarium. I am grateful have had access to the decades-worth of specimens collected by the Auckland Botanical Society. Lastly, I thank J. Schmack for her ongoing love and support.

